# An AI Agent for cell-type specific brain computer interfaces

**DOI:** 10.1101/2025.09.11.675660

**Authors:** Arnau Marin-Llobet, Zuwan Lin, Jongmin Baek, Almir Aljovic, Xinhe Zhang, Ariel J. Lee, Wenbo Wang, Jaeyong Lee, Hao Shen, Yichun He, Na Li, Jia Liu

## Abstract

Decoding how specific neuronal subtypes contribute to brain function requires linking extracellular electrophysiological features to underlying molecular identities, yet reliable *in vivo* electrophysiological signal classification remains a major challenge for neuroscience and clinical brain-computer interfaces (BCI). Here, we show that pretrained, general-purpose vision-language models (VLMs) can be repurposed as few-shot learners to classify neuronal cell types directly from electrophysiological features, without task-specific fine-tuning. Validated against optogenetically tagged datasets, this approach enables robust and generalizable subtype inference with minimal supervision. Building on this capability, we developed the BCI AI Agent (BCI-Agent), an autonomous AI framework that integrates vision-based cell-type inference, stable neuron tracking, and automated molecular atlas validation with real-time literature synthesis. BCI-Agent addresses three critical challenges for *in vivo* electrophysiology: (1) accurate, training-free cell-type classification; (2) automated cross-validation of predictions using molecular atlas references and peer-reviewed literature; and (3) embedding molecular identities within stable, low-dimensional neural manifolds for dynamic decoding. In rodent motor-learning tasks, BCI-Agent revealed stable, cell-type-specific neural trajectories across time that uncover previously inaccessible dimensions of neural computation. Additionally, when applied to human Neuropixels recordings–where direct ground-truth labeling is inherently unavailable–BCI-Agent inferred neuronal subtypes and validated them through integration with human single-cell atlases and literature. By enabling scalable, cell-type-specific inference of *in vivo* electrophysiology, BCI-Agent provides a new approach for dissecting the contributions of distinct neuronal populations to brain function and dysfunction.

## Main

Decoding neural activity with spatial, temporal and cell-type specificity remain a fundamental and pressing challenge in systems neuroscience and brain-computer interface (BCI) research^1^. High-density recordings using rigid Neuropixels probes^2–4^ or flexible electrodes^5–8^ provide single-cell resolution in behaving animals but rely heavily on subject-specific decoders and lack molecular cell-type specificity^9^ (**Fig. 1a**). Techniques for precise molecular cell-type identification, including optogenetic^10^ and chemogenetic^11^ tagging methods, offer specificity but suffer from low throughput in terms of the number of identifiable cells and limited chronic stability. Furthermore, despite the availability of comprehensive single-cell molecular and spatial transcriptomic atlases^12, 13^ and extensive literature^14^ detailing cell-type-specific properties, current methodologies lack the capacity to integrate this rich information with real-time *in vivo* electrophysiological recordings. Achieving such integration is essential, as understanding neural computations at the cell-type level would greatly enhance the interpretability of neural data, revolutionize therapeutic strategies by enabling the tracking of specific neuronal populations in neurological disorders, and promote the development of highly precise neural interfaces.

**Fig 1:**
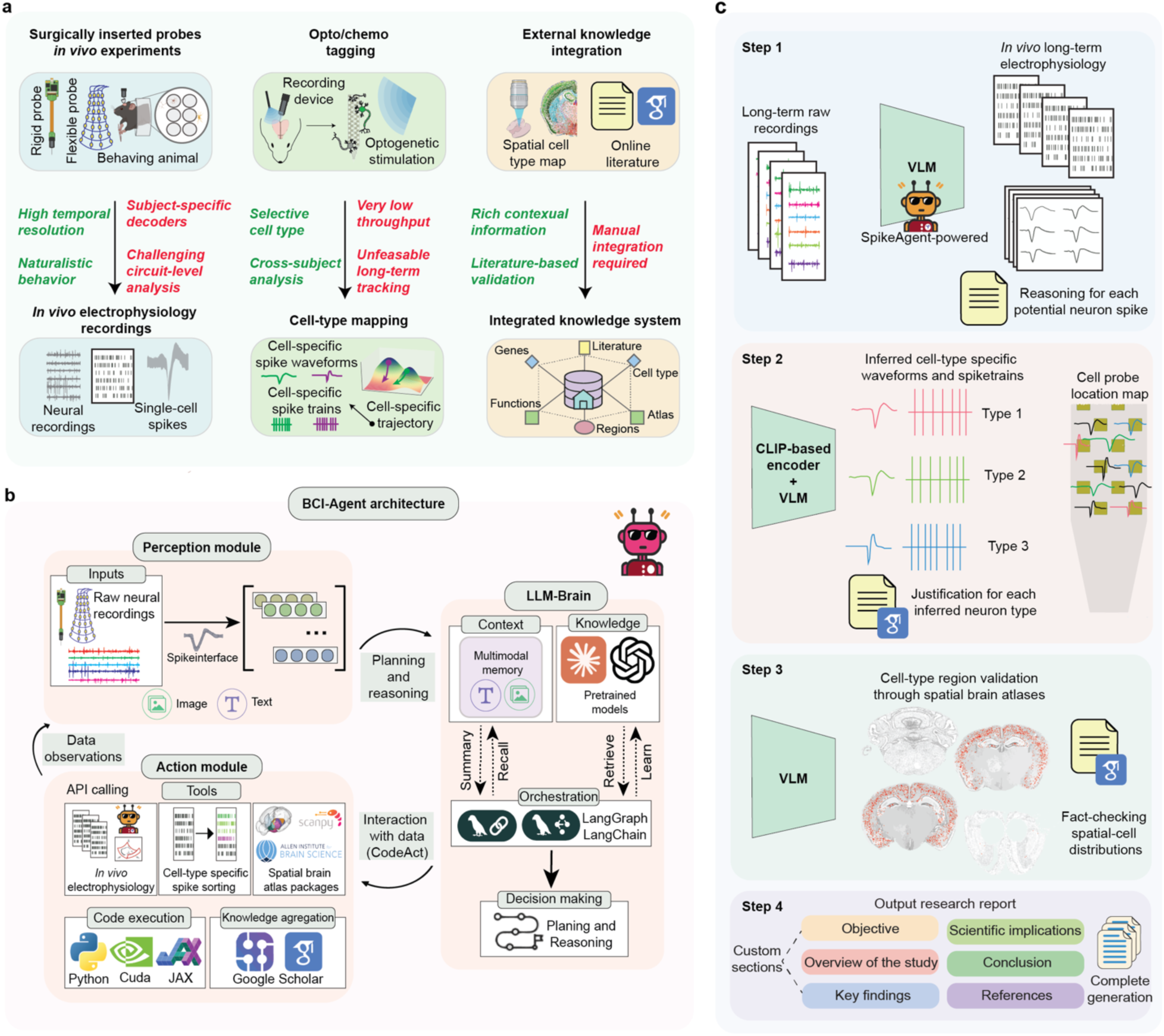
Overview of brain computer interface AI agent. a, Current methodologies for analyzing *in vivo* electrophysiological recordings. Surgically inserted probes capture neural activity at high temporal and single-cell resolutions during naturalistic behaviors but require subject-specific decoders, complicating circuit-level analysis. Optotagging and chemotagging provide selective cell-type and cross-subject analyses but suffer from low throughput and is impractical for long-term tracking. External knowledge integration through spatial cell-type atlases and literature offers contextual validation but relies heavily on manual, labor-intensive processes. **b**, System architecture of BCI-Agent, structured around perception, action, and language reasoning modules. The Perception module processes raw neural recordings, translating them into visual and textual features. These are passed to the LLM-Brain, a vision-language-powered module integrating multimodal memory, pretrained models, and literature retrieval for context-aware reasoning. The Action module executes decisions using a variety of tools: spike sorting pipelines, spatial brain atlas queries, and electrophysiology tools. It interacts with the data through API calls, code execution (Python, CUDA, JAX), and literature-based knowledge aggregation (Google Scholar). **c**, Processed data workflow of BCI-Agent for long-term *in vivo* electrophysiology. Raw neural recordings across days are fed into a VLM-powered spike sorting module pipeline. Then, each spike waveform and other neuron features are processed through a CLIP-based encoder to infer cell types, with accompanying reasoning and justifications. Finally, the inferred neurons are spatially mapped to anatomical probe locations and validated using reference spatial transcriptomic atlases, enabling biologically grounded interpretations of neural population activity across time. All agent-generated justifications and references are archived for future use. The dataset and its agentic results are compiled into structured, scientifically interpretable research reports including objectives, methods, findings, conclusions, and referenced biological literature, ensuring comprehensive archiving for future analysis.

Traditional machine learning approaches have attempted cell-type classification using electrophysiological features^15–17^ such as waveform templates, inter-spike intervals (ISI), and autocorrelograms (ACG), but face fundamental limitations. These methods depend on large, expert labeled datasets and rigid input-output mappings, limiting their generalizability across diverse experimental conditions and brain regions. The systematic integration of external resources, such as molecular atlases and scientific literature, remains unexplored, constraining validation of findings and hypothesis generation. Furthermore, although low-dimensional neural analyses have provided insights into neuron population dynamics^18, 19^, existing approaches require extensive manual intervention and rarely incorporate cell-type specificity. To overcome these limitations, we need methods that can infer cell types across diverse experimental conditions while simultaneously integrating existing molecular, literature-based knowledge and neural computation tools.

Recent advances in artificial intelligence (AI) offer promising approaches to these challenges. General purpose vision-language models (VLMs)^20–22^, particularly those using Contrastive Language-Image Pretraining (CLIP)^23^ such as OpenCLIP^24^ or SigLIP^25, 26^, have demonstrated remarkable abilities to extract meaningful patterns from visual data^27^. These models, trained on millions of image-text pairs, excel at few-shot learning–recognizing subtle visual distinctions without domain-specific training. More recently, the emergence of agentic AI^28–30^–systems that autonomously perceive, reason, plan, and execute complex workflows–has transformed scientific automation and discovery across fields including molecular biology^31^, chemistry^32^, behavior^33^, spatial biology^34^ and robotics^35, 36^. These agents orchestrate multiple tools, synthesize heterogeneous information, and embody scientific reasonings^37, 38^. We hypothesized that VLMs could decode cell-type signatures when electrophysiological features are transformed into a visual recognition task. More importantly, we recognized that by embedding this capability within an agentic framework, we could create a system that mirrors expert neuroscientists’ analytical workflows–not just classifying neurons, but validating predictions against molecular atlases, cross-referencing findings with peer-reviewed literature, and tracking their neural manifold dynamics to enable the study of how cell types contribute to behavior over extended timescales. This agentic orchestration would transform isolated cell-type classification into scalable, biologically grounded understanding and processing of neural recordings.

Here, we demonstrate that pretrained vision-language models can indeed classify neuronal cell types with remarkable accuracy. When electrophysiological features are converted to standardized images, recent models like SigLIP distinguish excitatory and inhibitory subtypes using just a few examples per type, without neuroscience-specific training. However, classification alone cannot address the complexity of neural analysis. We therefore developed Brain-Computer Interface AI Agent (BCI-Agent), an autonomous AI system that transforms this finding into comprehensive cell-type-aware neural decoding multi-agentic system. BCI-Agent orchestrates three integrated capabilities: (1) VLM-based inference that accurately identifies neuronal subtypes from electrophysiological features across recording modalities; (2) biological validation through automated integration of molecular atlases and peer-reviewed literature, ensuring predictions align with established knowledge; and (3) incorporation of cell-type identity into long-term low-dimensional neural dynamics analysis, revealing how distinct populations contribute to behavior over extended timescales. This framework enables, for the first time, molecularly informed analysis of chronic recordings.

We validated BCI-Agent across three datasets: optogenetically-tagged ground truth rodent recordings from diverse brain regions, unlabeled chronic rodent recordings in visual and motor learning tasks, and human Neuropixels data. For experiments lacking direct validation, BCI-Agent identified cell types, linked these to molecular identities through atlas comparison and literature support, and generated cell-type-specific trajectories across time and subjects. Specifically, we leveraged BCI-Agent to study cell-type specific manifolds during long-term motor learning recordings, revealing distinct contributions of different neuronal populations to behavioral adaptation. In human recordings where ground-truth validation is inherently unobtainable, BCI-Agent provided biologically plausible cell-type assignments validated through cross-referencing with single-cell atlases and existing literature. BCI-Agent enables scalable cell-type inference in *in vivo* electrophysiology. It automatically adapts to species-specific atlases, synthesizes relevant literature, and generates testable hypotheses. Through these capabilities, BCI-Agent addresses fundamental challenges in systems neuroscience opening immediate applications from identifying cell-type-specific dysfunction in disease to potentially designing targeted neuromodulation therapies.

## Results

### BCI-Agent system architecture

BCI-Agent is composed of three core modules, Perception, Reasoning, and Action, which work in an iterative, agentic loop to autonomously analyze neural data (**Fig. 1b**). The Perception module is specialized in processing raw electrophysiological recordings and any associated contextual metadata. Using interfaces such as Spikeinterface^39^, this module converts raw signals into structured multimodal representations comprising image, text, and metadata that capture relevant features of neural activity. These representations are then passed to the Reasoning module, referred to as LLM-Brain, which directs the downstream analysis. The LLM-Brain is the core of the BCI-Agent, which employs LLMs such as Claude-3.7-sonnet^40^, GPT-4o^41^, and Gemini Pro-2.5^42^ as well as VLMs from the CLIP family such as Open-CLIP^24^ and SigLIP^25, 26^. These models leverage visual encoders to enable context-aware cell-type classification. Additionally, LLM-Brain manages multimodal memory, synthesizes information across modalities, and generates structured reasoning outputs. The Action module executes computational tasks guided by LLM-Brain, such as cell-type-specific spike sorting, spatial mapping using brain atlases, and validation through scientific literature queries via APIs. All computational operations are conducted within a traceable Python environment optimized with Compute Unified Device Architecture (CUDA) and JAX for efficient processing and scalability. Importantly, BCI-Agent’s iterative design ensures continuous refinement; outputs from the Reasoning module trigger computational tasks within the Action module, whose results feed back into Perception, enabling dynamic and adaptive analysis cycles.

A key structural element of BCI-Agent is the Code as Action (CodeAct) framework^43^, implemented across both the Reasoning and Action modules. CodeAct enables the autonomous generation and orchestration of the dynamic workflow by the agent and execution of code for each task. Meanwhile, multimodal memory retains brain probe metadata, intermediate data visualizations, user inputs, and analytical outputs (**Extended Data Fig. 1**), allowing the agent to maintain context over time and dynamically refine its reasoning, understanding and actions. Task orchestration, planning, and retrieval of external resources are managed via LangGraph and LangChain^44^, which facilitate memory recall, integration of external tools and autonomous decision-making during the process with minimal supervision.

In practice, when raw inputs of electrophysiological data and behavioral contexts are received, BCI-Agent autonomously invokes all three modules in sequence for an end-to-end data analysis. Raw inputs are processed through spike sorting, cell-type inference, and biological validation, integrating analytical tools with external databases to yield biologically meaningful results as outputs. The entire process is autonomous without manual intervention, and all the decision and outputs are traceable, saved as structured formats (e.g., CSV or JSON).

Importantly, this end-to-end analysis enables few-shot learning for neuronal subtype inferring. The VLM-driven spike sorting tracks signals from the same neurons across different sessions (**Fig. 1c**, Step 1). Extracted spike features are then converted into images and analyzed using CLIP-based encoders for few-shot neuronal subtype classification (**Fig. 1c**, Step 2). Inferred cell types are localized according to the region of implantation and recording metadata and validated via spatially resolved brain atlas data and relevant literature (**Fig. 1c**, Step 3). Finally, results are compiled into structured reports with all the decision-making process documented for transparency and reproducibility (**Fig. 1c**, Step 4). We demonstrated BCI-Agent’s capabilities through the following three applications: (1) automated multi-session neural data analysis, (2) cell-type inference through few-shot learning, and (3) external database integration for data validation.

### General-purpose contrastive pretrained visual language models are few-shot cell-type learners

Achieving molecular cell-type specificity from electrophysiology, regardless of brain region or recording technology, remains a fundamental challenge. While optogenetic tagging provides ground truth, it is limited to one or two cell types per experiment. Recent breakthroughs in computer vision and robotics have demonstrated that contrastive learning models, particularly CLIP (Contrastive Language-Image Pre-training), excel at extracting meaningful patterns from visual data across diverse domains^27^. These models learn robust visual representations by training on millions of image-text pairs, developing the ability to distinguish subtle visual features. We hypothesized that by converting electrophysiological time series, such as spike waveforms, ISI, and ACG, into standardized visual representations, we could leverage these powerful pretrained models to decode cell-type signatures that are challenging to capture in conventional models.

To test this hypothesis, we developed a few-shot classification framework using frozen pretrained CLIP encoders, specifically focusing on SigLIP^25^, one of the most recent CLIP-based vision-language models that employs advanced contrastive learning techniques (**Fig. 2a**). Importantly, in this approach, electrophysiological features from *K* labeled examples per cell type (support set) and unlabeled query neurons were converted to images and processed through the visual encoder. Classification occurred by computing distances between query embeddings and class prototypes formed from the support embeddings. We evaluated performance on the CellExplorer dataset⁴⁰, containing optotagged-validated neurons from different brain regions and across cell types such as juxtacellular neurons, pyramidal cells, SST interneurons, and others (**Extended Data Fig. 2a**).

**Fig 2:**
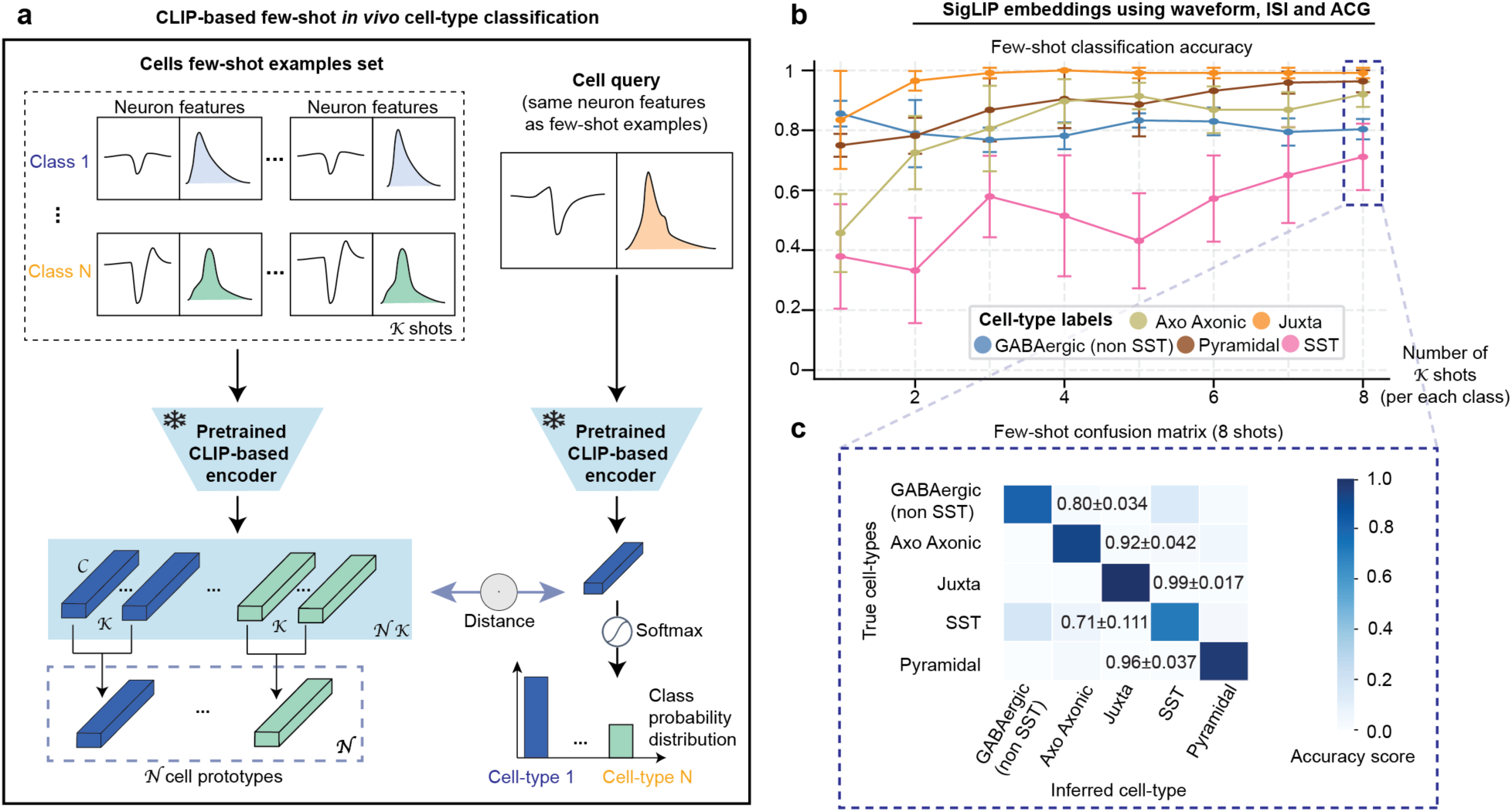
Vision-language models enable few-shot classification of *in vivo* neuron cell types from electrophysiological features images. **a,** Schematic of a CLIP-based few-shot classification framework using a frozen pretrained SigLIP image encoder for *in vivo* neuron cell-type classification. Both the support set and query image are embedded, class prototypes are formed, and classification is based on distance to prototypes. **b,** Few-shot classification accuracy using the SigLIP encoder across increasing numbers of shots. **c,** Confusion matrix for the SigLIP-based model using 8-shot classification. All results are computed over the same five random seeds using the same visual representations (waveform, ISI, ACG) and few-shot examples set for direct comparison.

SigLIP achieved robust few-shot classification, with accuracies above 70% across all cell-type categories and an average of 87% using only eight labeled examples per class (**Fig. 2b**). Without any neuroscience-specific training or fine-tuning, the model reliably distinguished excitatory pyramidal neurons from inhibitory subtypes, with misclassification rates below 5%. Most errors occurred within the GABAergic interneuron group, where however classification remained consistent, reaching ∼80% for non-SST and ∼71% for SST interneurons (**Fig. 2c**).

To validate that visual encoding was essential for this performance, we tested the same few-shot framework using raw time series features as embeddings without image conversion (**Extended Data Fig. 2b**). Notably, this approach can only achieve less than 40% accuracy even with eight examples for training per class. This notable contrast indicates that the pretrained visual representations are crucial for capturing cell-type distinctions. We also evaluated OpenCLIP^24^, another CLIP variant, and systematically tested different feature combinations by training random forest classifiers on the generated embeddings (**Extended Data Fig. 2c**). Both SigLIP and OpenCLIP maintained high accuracy (∼80%) across all feature combinations. UMAP visualizations revealed clear cell-type clustering in the embedding space, particularly for SigLIP, further suggesting that these models learn biologically meaningful representations without any neuroscience-specific training (**Extended Data Fig. 2d**).

These findings confirm that contrastive VLMs, originally trained on natural images and non-neuroscience specific, can also capture electrophysiological signatures that define neuronal cell types. The ability to achieve accurate classification with few-shot examples suggests this approach could generalize to diverse experimental contexts where obtaining larger labeled datasets is challenging. This discovery provides the foundation for BCI-Agent, enabling automated cell-type inference from standard electrophysiological recordings and opening new possibilities for understanding and capturing cell-type-specific computation *in vivo*.

### Cell-type reasoning grounding of predictions through anatomical and molecular context

While CLIP-based models excel at few-shot classification, real-world electrophysiological experiments rarely have optogenetic ground-truth labels. Therefore, ensuring the biological plausibility and interpretability of inferred neuron types is critical: predictions must align with known molecular markers, anatomical distributions, and established literature. To bridge this gap, we designed an integrated reasoning framework within BCI-Agent that automatically grounds cell-type predictions in existing neuroscience knowledge (**Fig. 3a**). This multi-stage validation operates autonomously: (1) initial classification using pretrained CLIP models, marking uncertain cases as “Unknown”; (2) detailed justifications generated by VLM instances that compare query features against reference patterns, providing interpretable explanations; and (3) biological validation through cross-referencing spatial transcriptomic atlases and automated literature synthesis, ensuring predictions match known molecular and anatomical properties.

**Fig 3:**
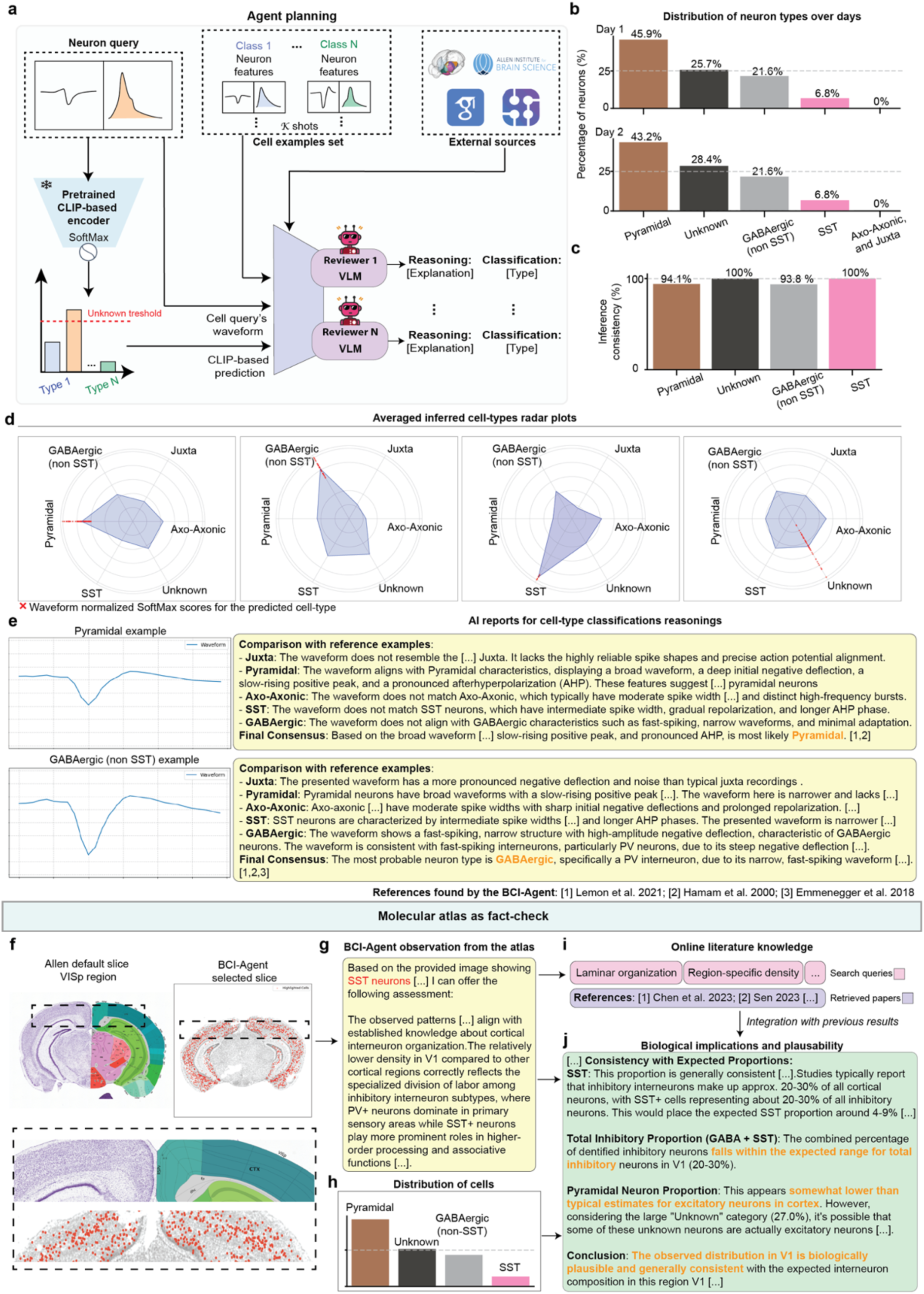
Integrated framework for biologically grounded cell-type classification and validation using BCI-Agent. **a**, Schematic of the BCI-Agent cell-type classification and reasoning pipeline. A waveform query is passed through a CLIP-based classifier to generate an initial cell-type prediction. This prediction is compared against a support set of example waveforms for known classes. Multiple VLM “reviewers” independently evaluate the query waveform by referencing known waveform features and literature, producing classification decisions and natural language justifications. The framework includes access to external literature databases (i.e. Allen Brain Atlas) for evidence retrieval. **b**, Distribution of predicted neuron types across two recording days. Bottom: c, Cross-day consistency comparison for inferred cell-type labels. **d**, Radar plots representing the similarity scores between the query waveform and different cell-type categories, as assessed by the VLM reviewers. Red X marks indicate the predicted class for the query waveform. **e**, Examples of VLM-generated classification reports. Each panel includes comparisons between the query waveform and reference waveforms across different neuron types. Reviewer reasoning and final consensus labels are generated with references to known electrophysiological features and classification heuristics based on prior studies^59–61^. **f**, Coronal brain slices showing the default reference section from the Allen Brain Atlas (left) and the most representative slice selected by BCI-Agent for the target VISp region (right). **g**, Text summary of BCI-Agent’s visual observations derived from the selected brain atlas slice. **h**, Bar plot showing the distribution of predicted neuron types based on waveform classification as part of the inputs of the data integration step for literature retrieval for validation. **i**, Literature retrieval component displaying example queries and retrieved references related to region-specific cell-type organization and laminar structure^12, 62^ **j**, Summary box integrating predictions with retrieved literature and molecular atlas-based expectations, used to assess the biological plausibility of inferred cell-type distributions.

To validate whether this framework produces biologically meaningful results without ground truth, we applied BCI-Agent to unlabeled chronic Neuropixels recordings from primary visual cortex (V1) collected over two consecutive days. Notably, BCI-Agent performed the entire pipeline autonomously–from spike sorting across sessions (**Extended Data Fig. 3**) to final cell-type assignment with molecular validation. Our evaluation examined three critical capabilities: (1) whether classifications remained consistent across days and aligned with expected V1 cell-type distributions, (2) whether the system could generate interpretable, literature-grounded justifications for each prediction, and (3) whether molecular atlas queries confirmed appropriate marker expression for the predicted cell types in the recorded anatomical location.

Initial waveform-based classifications revealed that the primary neuronal subtypes including Pyramidal, GABAergic (Non-SST), SST, and Unknown, were consistently identified across recording sessions (**Fig. 3b**). High agreement (above 93% in all cell-type classes) in classification across days (**Fig. 3c**) suggest the stability and internal consistency of the classification pipeline. Further validation using radar plots demonstrated that inferred inhibitory neuron classes (GABAergic and SST) exhibited waveform features distinct from excitatory neurons, reinforcing confidence in subtype assignments through the normalized SoftMax scores (**Fig. 3d**).

Next, we applied the VLM-powered reasoning module, which provided transparent explanations and literature-based support for each cell-type assignment. For instance, the VLM module of the BCI-Agent identified features such as “*broad waveform, deep initial negative deflection, slow-rising positive peak, and pronounced afterhyperpolarization*” as indicative of pyramidal neurons, and “*steep negative deflection and narrow, fast-spiking waveform*” characteristic of GABAergic neurons (**Fig. 3e**). Each explanation was explicitly linked to relevant literature, ensuring traceable and scientifically grounded reasoning and mitigating potential errors from pretrained model inference.

To evaluate biological plausibility, BCI-Agent incorporated spatial and molecular validation using the Allen Institute’s mouse brain atlas. The agent can autonomously identify the most representative slice for SST expression within V1 (**Extended Data Fig. 4b**). Leveraging this spatial context, VLM analyzed regional SST expression, providing structured biological insights consistent with established cortical interneuron distributions (**Fig. 3f-h**). The observed sparse SST expression in V1 matched known cortical patterns, wherein PV neurons typically dominate primary sensory areas while SST neurons are relatively more common in associative areas. This anatomical reasoning was further reinforced by automated literature integration (**Fig. 3i**).

Based on inferred neuronal subtype distributions (**Fig. 3g**), BCI-Agent generated a comprehensive biological plausibility report (**Fig. 3j**). The analysis confirmed that the inhibitory neuron proportions aligned with expected cortical ranges, specifically noting that SST neurons fell within the typical 4-9% range from transcriptomic studies. Although pyramidal neuron proportions appeared lower than standard estimates, the report suggested that some neurons classified as “Unknown” might include atypical excitatory cells, reconciling the observed excitatory-inhibitory balance with established biological norms.

We also challenged the BCI-Agent’s ability to generate those plausibility reports in a control experiment. We intentionally altered the metadata to incorrectly indicate the recording location as the cerebellum rather than the cortex (**Extended Data Fig. 5**). The result of this control experiment showed that BCI-Agent detected critical inconsistencies, such as the improbable high proportion of pyramidal neurons and excessive “Unknown” classifications, highlighting significant anatomical misclassification. Additionally, the presence and proportions of GABAergic and SST-expressing neurons diverged from established cerebellar neuron populations, emphasizing the need for methodological refinement to ensure accurate biological alignment and interpretability.

Overall, by combining waveform-based classification with interpretable VLM-based reasoning, real-time literature search and molecular atlas validation, represents a significant advance in neural computation. By providing multiple layers of evidence and validation, BCI-Agent transforms raw spike waveforms into biologically meaningful cell-type assignments with transparent justifications. This capability is particularly valuable for chronic recording experiments where direct molecular identification is impractical to be obtained, enabling researchers to track cell-type-specific neural dynamics over extended periods with higher confidence in the biological identity of recorded neurons.

### Cell-type-specific neural dynamics during motor learning

Having validated BCI-Agent’s ability to accurately classify and biologically ground cell types, we next tested whether these capabilities could reveal cell-type-specific dynamics during complex behaviors. By combining all the previously described modules, BCI-Agent offers a cohesive approach for understanding how different neuronal cell types uniquely contribute to behavior across long-term recordings (**Fig. 4a**). We tested this framework using 50 days of motor-learning data obtained from stable, flexible, high-density electrodes^45^. BCI-Agent successfully identified and stably tracked neuronal subtypes throughout the experiment, providing consistent and stable tracking. Additionally, it integrated these results with existing literature to offer deeper biological context and insights into the dynamics of neural activity.

**Fig 4:**
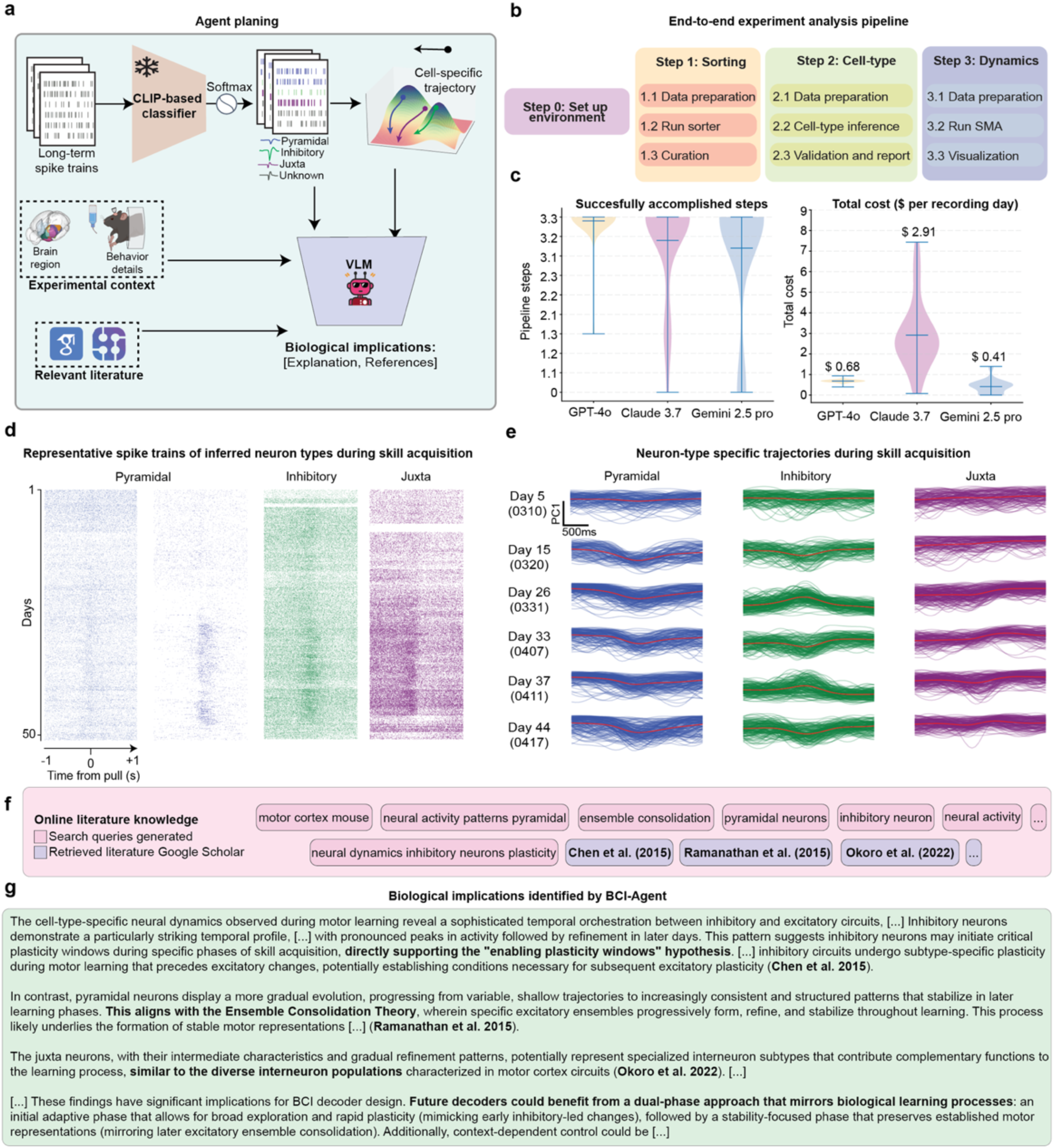
BCI-Agent enables long-term cell-type-specific neural manifolds during skill acquisition. **a,** Overview of the BCI-Agent analysis pipeline. Long-term spike trains are processed by a pretrained CLIP-based classifier to infer putative general neuron types (Pyramidal, Inhibitory, Juxta, or Unknown). These outputs, alongside experimental metadata (brain region, behavioral epochs), are passed to a VLM that generates structured explanations, retrieves relevant literature, and infers biological implications**. b**, End-to-end experimental pipeline, detailing the sequential steps from environment setup through spike sorting, neuron-type inference, validation, and dimensionality reduction neural dynamics. **c**, Benchmark of successfully accomplished steps without code failure and total cost usage, comparing the reliability of each LLM backend in executing tasks for a single recording day. **d,** Representative spike trains for each inferred neuron type recorded across consecutive days of motor skill learning. e, Visualization of low-dimensional neuron-type-specific population trajectories across multiple days of skill acquisition. Rows indicate recording days; columns show distinct neuron subtypes. Each plot visualizes smoothed, low-dimensional trajectories of neural activity over time. **f,** Online literature integration module. Automatically generated search queries (top) retrieve peer-reviewed publications^47, 48, 50^. **g,** Biological insights extracted by the system grounding observations in existing knowledge. All violin plots represent the distribution of each data over different seeds (N=30), with width indicating frequency. Horizontal lines show the mean, maximum, and minimum.

The analysis began with BCI-Agent autonomously performing spike sorting on the initial recordings to generate high-confidence neuronal labels (**Extended Data Fig. 6a-c)**. These labels were then used to train a lightweight, agent-supervised decoder that robustly tracked the same neurons in real-time across the subsequent weeks of the experiment (**Fig. 4b**, **Extended Data Fig. 6d-f**). With stable, long-term tracking established, the agent classified the spike trains into broad neuronal types (e.g., pyramidal, inhibitory, juxtacellular) and visualized their respective spiking activity patterns (**Fig. 4d**). We took this approach, combining inhibitory subtypes like GABAergic and SST into one category, due to the limited count of recorded neurons. The population dynamics for each cell type were then embedded into low-dimensional manifolds using an adapted version of Similarity Matching Algorithm (SMA), producing neural trajectories that evolved as the animals acquired the motor skill (**Fig. 4e**). We confirmed the robustness of this end-to-end pipeline by benchmarking it with our reasoning models, including GPT-4o, Claude 3.7, and Gemini 2.5-Pro, all of which successfully executed the required analytical steps. The entire analysis was also highly efficient, costing less than one dollar per animal per recording day (**Fig. 4c**).

Beyond visualization, BCI-Agent employs a structured, three-layer scientific reasoning approach (**Extended Data Fig. 7**): (1) internal neuroscience knowledge retrieval via large language models, (2) local trajectory analysis using VLMs to detect critical patterns (e.g., trajectory valleys, U-shaped dynamics), and (3) external validation using automated literature retrieval. This comprehensive analytical framework allowed BCI-Agent to autonomously generate structured, natural-language scientific reports detailing how cell-type-specific neural dynamics evolved during motor learning. Reports included summaries of experimental overviews, references to relevant literature, discussions of biological implications, and transparent acknowledgments of methodological limitations (**Fig. 4f-h**).

For example, BCI-Agent identified clear stabilization trajectories in pyramidal neurons, transitioning from broad exploratory activity to compact, task-specific manifolds, consistent with the formation of specialized excitatory ensembles. In contrast, inhibitory neurons showed earlier, more variable shifts aligning with their hypothesized role in initiating plasticity and modulating network dynamics during early learning phases. Juxta neurons exhibited intermediate behaviors, stabilizing later and seemingly mediating interactions between excitatory and inhibitory populations.

Crucially, every hypothesis produced by BCI-Agent is both data-driven and literature-grounded. For instance, the observed U-shaped activity in pyramidal neurons is not merely descriptive; it is contextualized within literature on skill acquisition that highlights sequential phases of exploration, plasticity, and stabilization^46–48^. Similarly, interpretations of inhibitory neuron behavior are anchored in studies on plasticity gating, noise regulation, and inhibitory remapping. These insights align with theoretical frameworks of ensemble consolidation and cortical plasticity gating^47, 49, 50^. Notably, BCI-Agent also identified *Chen et al.* (2015)^47^, *Ramanathan et al.* (2015)^48^, and *Okoro et al.* (2022)^50^ as the key works that link inhibitory circuit dynamics to the stabilization of excitatory ensembles and the emergence of sparse, task-tuned representations during learning. These citations are not hallucinated; they are retrieved, validated, and explicitly referenced through a transparent, reproducible pipeline by the agent.

Ultimately, BCI-Agent represents a novel paradigm from passive data analysis towards active scientific exploration and hypothesis-driven interpretation. Its end-to-end reasoning, from raw spike data to biologically plausible insights, demonstrates the substantial potential of agentic AI to serve as a co-scientist, providing scalable, reproducible, and interpretable solutions essential for advancing systems neuroscience and next-generation brain-computer interfaces.

### BCI-Agent translates cell-type inference methods to human electrophysiology

The ultimate test of BCI-Agent’s generalizability lies in human applications, where conducting molecular cell-type identification experiments has remained challenge due to technical and ethical constraints. However, recent comparative studies have demonstrated conserved cellular architectures between rodents and primates, supporting cross-species neuron-type inference. For example, successful cell-type classification across cerebellar regions in mice and non-human primates revealed conserved population dynamics^9^, while detailed single-nucleus RNA-sequencing analyses highlighted similarities and key species-specific differences between human and rodent cortical neuron populations^51^. Collectively, these findings suggest that electrophysiological signatures used by BCI-Agent for rodent neuron classification may generalize effectively to human recordings.

To test this possibility, we applied the BCI-Agent framework to human Neuropixels recordings from the motor cortex of a human subject undergoing awake surgical resection^52^. Using pretrained SigLIP embeddings derived from rodent ground-truth waveforms (**Fig. 5a**), we classified human neuronal waveforms, achieving meaningful cross-species correspondence in neuron embeddings (**Fig. 5b**). Radar plots further confirmed consistent electrophysiological signatures across inferred human neuronal types, analogous to rodent classes (**Fig. 5c**). Crucially, BCI-Agent’s inferred cell-type labels allowed us to extract human-specific measurements, such as cell-type-specific firing rates (**Fig. 5d**), providing potentially detailed insights into neuronal subtype physiology in the human cortex, previously challenging to obtain.

**Fig. 5:**
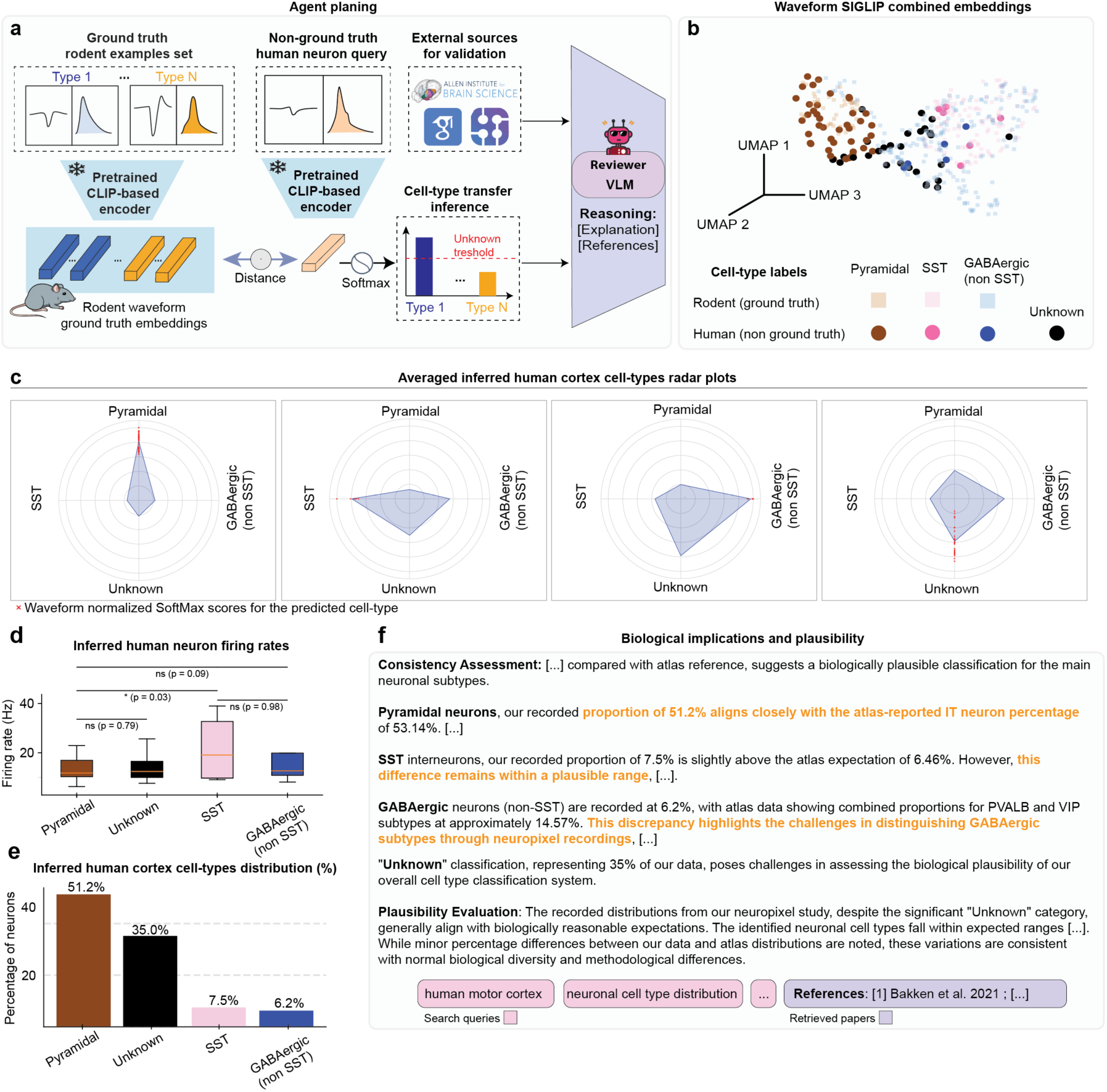
Cross-species transfer of neuronal cell-type inference from rodent to human single-neuron recordings in motor cortex. **a,** Schematic overview of the cell-type inference and validation pipeline. Rodent neuronal waveforms with known ground-truth labels are encoded using a pretrained CLIP-based model to create reference embeddings. Human neuron waveforms from Neuropixels recordings (without ground-truth labels) are similarly encoded and classified by comparing their embeddings to the rodent reference set. A VLM integrates external sources to validate and provide biological reasoning for inferred cell-type assignments. **b**, UMAP visualization of combined rodent (ground truth) and human (non-ground truth) neuronal waveform embeddings generated by the pretrained CLIP-based model, illustrating the clustering of inferred cell types. **c**, Radar plots showing the waveform normalized SoftMax scores for the inferred human cortical neuron types. **d**, Box plots of inferred human neuron firing rates, comparing differences among identified neuronal subtypes. **e**, Bar plot summarizing the inferred neuronal cell-type distribution percentages from human Neuropixels recordings, identifying proportions of pyramidal, SST, GABAergic (non-SST), and unknown neuron classes. **f**, Summary of biological plausibility and consistency assessment comparing inferred human neuronal subtype proportions against established reference data from molecular atlases and literature^63^. All box plots represent median values (center lines), interquartile ranges (boxes), and 1.5× interquartile range whiskers. Statistical significance indicated by asterisks: *P < 0.05, **P < 0.01, and ns = not significant.

Quantitative analysis of inferred human neuronal types revealed predominantly pyramidal neurons (51.2%), followed by a substantial “Unknown” category (35%), with smaller populations of SST interneurons (7.5%) and GABAergic non-SST interneurons (6.2%) (**Fig. 5e**). Biological plausibility assessments via automated literature retrieval and leveraging a single-cell human atlas^53^ comparisons suggested that the inferred neuronal proportions closely matched human cortical distributions reported in molecular atlas studies, validating our approach (**Fig. 5f**).

We further applied the same framework to another human Neuropixels’ recording from the dorsolateral prefrontal cortex^54^ (DLPFC) and found similar proportions and inferred results (**Extended Data Fig. 8a-b).** Although this region is not included in the molecular single-cell dataset used for validation, BCI-Agent autonomously identified and compared it to the closest available region in the dataset (medial prefrontal cortex), justifying this comparison as “*The closest region to the dorsolateral prefrontal cortex in the atlas is the medial prefrontal cortex which shares associative and cognitive functions and is functionally connected to the DLPFC*” (**Extended Data Fig. 8c**).

Overall, these results propose BCI-Agent as a novel potential method for cell-type inference to human electrophysiology, overcoming previous challenges by providing biologically meaningful, human-specific neuronal insights. At the same time, the approach has important limitations. Training on rodent ground-truth data may not capture the full diversity of primate or human neurons, and the absence of molecular validation in human recordings prevents definitive confirmation of the inferred classes. Future progress will depend on expanding ground-truth resources, increasing the use of diverse cell-types optotagging, and integrating electrophysiology with complementary modalities, with validation in non-human primates likely serving as an important step toward eventual clinical applications in human neuroscience.

## Discussion

BCI-Agent introduces a novel integrated approach to analyzing and interpreting large-scale neural data that offers new insights to rodent as well as human electrophysiology. Unlike traditional methods, which typically address isolated tasks such as signal extraction or dimensionality reduction independently, BCI-Agent integrates spike sorting, cell-type classification, neural manifold computation, and biological reasoning within a single, autonomous framework. This integration allows the system not only to process *in vivo* neural recordings but also to interpret them through established neuroscience theories, community resources and validated peer-reviewed literature. In this role, BCI-Agent functions effectively as an AI co-scientist^37, 38, 55^, designed explicitly to support and augment human researchers. Leveraging agentic AI, BCI-Agent generates rich electrophysiological datasets that combine precise spatiotemporal neural dynamics with inferred molecular annotations, thus facilitating biologically meaningful insights that previously were unrealistic to obtain at scale.

A core finding is the intrinsic capability of pretrained VLMs, particularly models like SigLIP, to differentiate neuronal cell types effectively through visual embeddings, even without explicit neuroscience-specific training. This discovery effectively reframes *in vivo* cell-type classification as an image recognition task, which can be addressed by utilizing pretrained VLMs originally developed for general-purpose feature extraction in computer vision. The inherent ability of these models to distinguish cell-type-specific visual patterns enables a new way of producing cell-type labels, reducing the reliance on extensive labeled datasets, enabling powerful few-shot classification.

On the technical side, BCI-Agent addresses three interconnected challenges that have limited progress in systems neuroscience. First, it achieves autonomous cell-type classification validated through molecular atlases and spatial transcriptomics, ensuring biological plausibility. Second, it systematically integrates external resources, such as peer-reviewed literature and molecular databases, for hypothesis generation and validation, preventing hallucinated interpretations. Third, it enables multi-session tracking of cell-type-specific dynamics over weeks to months, revealing how distinct neuronal populations contribute differentially to behavior. These capabilities emerge from BCI-Agent’s agentic architecture: autonomous task planning, multimodal memory maintenance, and iterative validation through external resources.

Our work underscores how well-orchestrated agentic systems uniquely enable seamlessly merging multiple disciplines and data modalities within a unified framework. BCI-Agent exemplifies this integrative approach, highlighting how agentic AI systems can serve as scientific reasoning engines capable of observing, contextualizing findings, and generating hypotheses in accessible human language. This capability presents significant opportunities for employing LLMs and VLMs as active collaborators rather than passive tools, contributing meaningfully to scientific exploration. Although BCI-Agent currently specializes in neural computational tasks, its foundational principles such as few-shot classification, multimodal memory, structured prompting, and literature-grounded reasoning are broadly applicable and have the potential to foster interdisciplinary breakthroughs across various biological and systems sciences.

Critically, the translation of BCI-Agent’s cell-type inference methodologies to human Neuropixels recordings enhances the translational and clinical potential of electrophysiological research. Historically, direct molecular identification of neuronal subtypes in human recordings has been unattainable due to ethical and technical constraints, severely restricting interpretability. By leveraging conserved electrophysiological signatures across species, as supported by comparative studies, BCI-Agent now enables inference of cell-type identities in human datasets. This advancement opens unprecedented avenues for exploring detailed neuronal characteristics in humans, such as cell-type-specific firing rates, substantially advancing our understanding of human cortical physiology. Consequently, BCI-Agent holds promise for providing biologically meaningful insights into neuronal dynamics, facilitating breakthroughs in both fundamental neuroscience and clinical applications.

Nonetheless, several significant challenges remain. First, the reliability of few-shot cell-type inference remains heavily dependent on the quality, diversity, and comprehensiveness of available reference datasets, which are currently limited mostly to cortical and hippocampal regions in rodents^17, 56^. Therefore, generalization to other brain regions in rodents as well as primates remains challenging and requires the expansion of reference datasets. Real-time integration with experimental hardware also requires significant advancement to achieve robust, high-throughput closed-loop *in vivo* applications. However, these limitations present clear innovation opportunities across both hardware and software domains. Recent advances in ultra-high-density recording technologies^57^ promise richer electrophysiological features for classification, while the rapid evolution of OpenCLIP to SigLIP demonstrates how quickly foundational models improve.. As foundational AI models continue to advance in accuracy, abstraction, and generalization capabilities, their application to few-shot and potentially zero-shot neural computation and multimodal reasoning will likely become even more robust and scalable^58^.

In conclusion, BCI-Agent offers neuroscientists a structured, interpretable, and reliable analytical framework tailored specifically for large-scale, multimodal, and continuous neural recordings. Traditional analysis approaches often fall short in providing the coherence, scalability, and clarity needed to effectively align long-term neural data with molecular identities, especially when reference data is sparse or limited. As neuroscience increasingly engages with complex and expansive datasets, frameworks like BCI-Agent will become indispensable. They enable researchers to efficiently navigate and integrate the extensive array of tools, databases, and knowledge available, ultimately making sense of the brain’s inherent complexity, variability, and rich behavioral dynamics.

## Methods

### BCI-Agent framework

We developed BCI-Agent using the LangChain and LangGraph AI agent frameworks to create an extensible, state-aware architecture for autonomous analysis of neural recordings. These frameworks support prompt management, tool invocation, memory handling, and conditional logic across multistep workflows. BCI-Agent leverages this infrastructure to perform tasks spanning from spike sorting and cell-type inference to neural decoding, spatial validation, and biologically grounded hypothesis and observation generation.

BCI-Agent incorporates advanced multimodal LLMs, including GPT-4o, Claude-3.7-Sonnet and Gemini 2.5-Pro, which were used in a zero-shot setting to retain their general-purpose reasoning capabilities. Vision-language capabilities are provided through OpenCLIP and SigLIP models for few-shot classification and anatomical image comparison. The architecture follows an adapted CodeAct-style (Code as Action) agent paradigm, enabling the agent to dynamically interpret multimodal inputs, retrieve context-specific knowledge, execute code, and iteratively refine analyses.

LangGraph’s cycle-aware logic underpins BCI-Agent’s iterative reasoning process, which unfolds through the following steps: (1) observation of local neural data (e.g., waveforms, spike trains, PCA trajectories), (2) multimodal reasoning and planning, (3) code generation and execution, (4) evaluation of outputs and contextual comparison to known biological patterns, and (5) integration of internal and external knowledge sources. Long-term coherence is maintained through explicit session state tracking and memory management, supporting extended, multi-turn scientific workflows. This includes integration with the Allen Institute’s Brain Atlas for spatial validation, SerpAPI ^64^and Google Scholar for real-time scientific literature retrieval, and multimodal data ingestion pipelines for spike waveforms and anatomical images.

### External reasoning integration

BCI-Agent includes a suite of modular tools inspired by the CodeAct architecture, enabling code-driven execution within a flexible reasoning environment. Tools span across data preprocessing, spike sorting (via SpikeInterface), unit visualization, and neural dynamics computation, each generating executable Python code tailored to the experimental context. A Python REPL interface allows BCI-Agent to dynamically interact with datasets, execute reasoning-informed curation decisions, and visualize results, all while tracking state and session history.

A key feature of BCI-Agent is its multimodal biological reasoning pipeline, which combines vision-language interpretation with external database grounding. For example, when classifying neuronal cell types, the agent validates predictions using the Allen Brain Atlas’s spatial expression maps, selecting representative brain slices, and evaluating anatomical similarity using VLMs. These inferences are then contextualized through LLM-generated explanations and checked against real-time literature using SerpAPI-powered scientific queries.

### Multimodal memory and neural context awareness

To support context-aware neuroscience reasoning, BCI-Agent employs a structured multimodal memory system. This memory persists in textual summaries, visual inputs (waveforms, ISI, ACG plots, PCA trajectories), intermediate outputs (e.g., spike sorting results, decoder weights), and user-agent interaction history. This enables the agent to interpret neural population dynamics in a temporal context, compare evolving patterns across sessions, and ground its insights into prior reasoning. The memory system also tracks classifier prototypes and spike feature embeddings over time, which is critical for long-term cell-type-specific tracking and population manifold reconstruction.

### Integrative tools backend core

#### Automated spike sorting and neural computation

For spike sorting, we adapted the SpikeAgent ^65^framework, an autonomous agent-based spike processing system designed for session-by-session sorting of neural recordings. SpikeAgent operates as a multimodal interface that combines code generation with structured reasoning to automatically process raw electrophysiological recordings. For each session, SpikeAgent uses built-in modules to handle data loading, bandpass filtering, common average referencing, spike detection, and sorting via SpikeInterface-compatible algorithm. For all our recordings we used Kilosort4^66^ for Neuropixels or Mountainsort4^67^ for other recordings.

Following spike sorting, the system performs automated unit curation using VLM reviewers, which assess waveform quality and inter-spike interval distributions to label units as “Good” or “Noise”. All steps, spike sorting, feature extraction, waveform visualization, and curation are fully automated and documented, enabling consistent processing across recording days.

#### Real-time spike classification with multi-feature neural decoder

To enable continuous spike classification in real-time or across future sessions, we implemented a lightweight neural decoder trained using output from the first SpikeAgent session. Our model was inspired by prior work employing multi-feature classification approaches^45^, but we used a single multi-class model that jointly distinguishes between noise and all neuron identities discovered on the initial day. The model’s output layer includes one “noise” class and one class for each sorted unit, detected in its training session. For all the subsequent sessions (after day or session 1) each detected spike, a multimodal feature vector is constructed using three input modalities: (1) the spike waveform snippet from the nearest channels, (2) waveform snippet from one extremum channel (3) spatial coordinates of extremum channel based on electrode layout. The model is trained using a categorical cross-entropy loss function, using labeled spikes from the initial day where neuron identities are known. Once trained, the decoder generalizes to future sessions, enabling real-time inference and consistent unit tracking without re-running spike sorting.

#### Neural dynamics extraction

To extract low-dimensional neural trajectories from spike trains, BCI-Agent uses a biologically inspired neural network that performs online dimensionality reduction in real time. This unsupervised neural network linearly maps each high-dimensional input vector of spike activity into a lower-dimensional space using a feedforward weight matrix. The weights are updated continuously using a Hebbian learning rule, where the matrix is adjusted at each time step based on the outer product of the output vector and the input, scaled by a learning rate. In parallel, a lateral inhibition matrix maintains competitive interactions between dimensions in the projected space, encouraging sparsity and orthogonality. To ensure smooth representation of neural population dynamics, the resulting low-dimensional trajectories are temporally filtered using a Gaussian convolution kernel.

### Few-shot cell-type classification with vision-language models

To classify neurons by putative cell type, BCI-Agent leverages pretrained CLIP-based VLM (Open-CLIP, SigLIP) in a few-shot learning framework. Spike waveform features including raw waveform plots, ISI histogram plots, and ACG images, were rendered as figures using default SpikeInterface color. These visual representations were embedded into the CLIP feature space, where cell-type classification was posed as a few-shot image recognition tasks.

For each cell type, *K* support examples were encoded to construct class prototypes. Query embeddings were then compared to these prototypes using cosine similarity, producing a probability distribution over candidate cell types. Importantly, no model fine-tuning or gradient updates were performed; the pretrained VLM was used in a frozen state.

We benchmarked classification performance across input modalities, comparing waveform-only inputs against composite representations (waveform + ISI; waveform + ISI + ACG). Cell-type labels include excitatory (pyramidal) neurons and multiple inhibitory neuron classes (PV, SST, VIP, VGAT, PVVGAT), along with Juxtacellular and Axo-axonic categories. Few-shot evaluations were conducted using 1 to 8 support examples per class, and all experiments were repeated across five random seeds to assess stability and variability.

### Anatomical and spatial validation

To validate the biological plausibility of inferred cell-type distributions, BCI-Agent performs anatomical context matching using VLMs to compare coronal brain slice images. A structured prompting system guides a multimodal VLM (Claude 3.7, GPT-4o, Gemini 2.5-Pro) to assess anatomical similarity between a slice focusing on any cell-type and a reference slice from the Allen Brain Atlas corresponding to a known brain region (e.g., primary visual cortex, V1). Each comparison is scored on a continuous scale (0.00 -1.00) based on structural features such as shape, landmark alignment, and laminar organization. A final similarity classification (“High”, “Good”, “Moderate”, “Low”) is assigned based on predefined thresholds in the system prompt.

To identify the most anatomically plausible slice for a given region, BCI-Agent loads all available slices with detectable expression of a target cell type and compares them to the canonical reference slice for that region from the Allen Brain Atlas. Each cell-type-positive slice is encoded as an image and submitted for pairwise comparison. Comparisons are run in parallel, and similarity scores are aggregated to determine the best-matching experimental slice.

This top-ranked slice is then used as a spatial anchor to contextualize cell-type distributions within known regional architectures. At this stage, BCI-Agent invokes its biological plausibility engine -a large language model system prompted with both the anatomical context and the inferred cell-type distribution. The system performs multi-level reasoning by combining: (1) prior knowledge of region-specific cell-type densities, (2) visual interpretation of laminar expression patterns, and (3) automated literature retrieval via SerpAPI and Google Scholar APIs. The final output includes a structured, natural language justification for the spatial plausibility of the observed distributions, supported by peer-reviewed references and confidence ratings.

### Biological synthesis and knowledge integration

BCI-Agent integrates biological knowledge by combining large-scale literature retrieval with internal neuroscience reasoning to interpret results across multiple modules. Once core analyses -such as cell-type classification, anatomical alignment, or neural trajectory extraction -are completed, the agent automatically initiates a knowledge synthesis phase. This phase uses a structured prompt system to engage LLMs with contextual cues from the data (e.g., spike dynamics, brain region, inferred cell types). The agent queries scientific search engines via SerpAPI and Google Scholar to retrieve peer-reviewed publications and combines these with its built-in neuroscience priors to generate biologically grounded explanations. These explanations include citations, summary statistics, and biologically plausible interpretations, allowing the system to validate its own outputs and generate hypotheses.

This synthesis layer is applied uniformly across several components of the framework: to assess the plausibility of inferred spatial expression patterns during atlas comparisons, to interpret cell-type-specific neural dynamics over time, and to ground predictions in known circuit functions. By fusing structured data with real-time literature and LLM-generated reasoning, BCI-Agent transforms local data plots into interpretable, evidence-backed insights.

### Electrophysiology data

All data used in this study was publicly available and required no additional animal experimentation. We analyzed two electrophysiological datasets collected from distinct rodents using different electrode technologies, a third reference dataset containing opto-tagged neuronal cell types, as well as a fourth human Neuropixels dataset.

#### Rodent Neuropixels dataset

The first dataset comprised raw extracellular recordings acquired with Neuropixels 2.0 probes^3^, which feature high-density site arrays and improved signal-to-noise characteristics. Data was obtained from a publicly released mouse visual cortex dataset. All experimental procedures, probe geometry, and behavioral protocols are documented in the original dataset publication. For our analysis, we focused on recordings from Subject AL031, specifically sessions “AL036_2020-02-19” and “AL036_2020-04-15” corresponding to day 1 and day 57 (or session 1 and 2) in our figures.

#### Flexible electrode dataset

The second dataset was derived from chronic motor cortex recordings using a flexible tetrode-like electrode array interfaced with an Intan RHD2132 amplifier for 50 almost consecutive days. Data acquisition was performed at 10 kHz using a custom PCB-based recording system. This dataset was originally published in conjunction with the flexible electrode and animal experiments characterization^45^.

#### CellExplorer opto-tagged reference dataset

To evaluate cell-type classification, we used the opto-tagged reference dataset provided by CellExplorer^56^, a MATLAB-based framework for standardized spike-based cell metrics and visualization. The dataset spans recordings from six laboratories and includes optogenetically identified inhibitory and excitatory neurons. We used spike waveform images, ISI histograms, and ACG exported via custom Python functions to serve as ground-truth-labeled examples for few-shot learning and benchmarking. CellExplorer’s pipeline supports multiple spike-sorting formats and includes extensive metadata, ensuring compatibility and interpretability in our analyses.

#### Human Neuropixels dataset

The last dataset includes two publicly available human Neuropixels recordings obtained during surgical procedures.

The first recordings were collected using standard Neuropixels probes during intraoperative monitoring associated with surgical resection. Complete documentation of experimental conditions, probe placement details, patient demographics, and clinical context are provided in the original dataset publication^52^.

The second recording was obtained from a patient undergoing deep brain stimulation (DBS) electrode placement, where the Neuropixels probe was inserted through the dorsolateral prefrontal cortex as part of the surgical trajectory. This recording provides complementary data from a different cortical region and surgical context. Complete documentation of experimental conditions, probe placement details, patient demographics, and clinical context are provided in the original dataset publication^54^.

## Code availability

Software code for this study will be made available at the time of publication at http://github.com/LiuLab-Bioelectronics-Harvard/BCI-Agent.

## Acknowledgements

A.ML. acknowledges the support from the RCC-Fellowship of Harvard University and the Excellence Fellowship of the Fundacion Rafael del Pino. We acknowledge support from NIH/NIDDK 1DP1DK130673 (J.Liu); NSF ECCS-2038603 (J.Liu and N.L.); and NIH/NLM 5R01LM014465 (J.Liu and N.L.).

## Author contributions

A.ML., Z.L., and J.Liu. conceived the study. A.ML. developed the methodology and conducted all experiments. A.ML. and J.B. prepared the figures. A.ML. and Z.L. drafted the manuscript. All authors provided critical feedback and contributed to the interpretation of results and figures.

---

**Extended Data Fig. 1.**
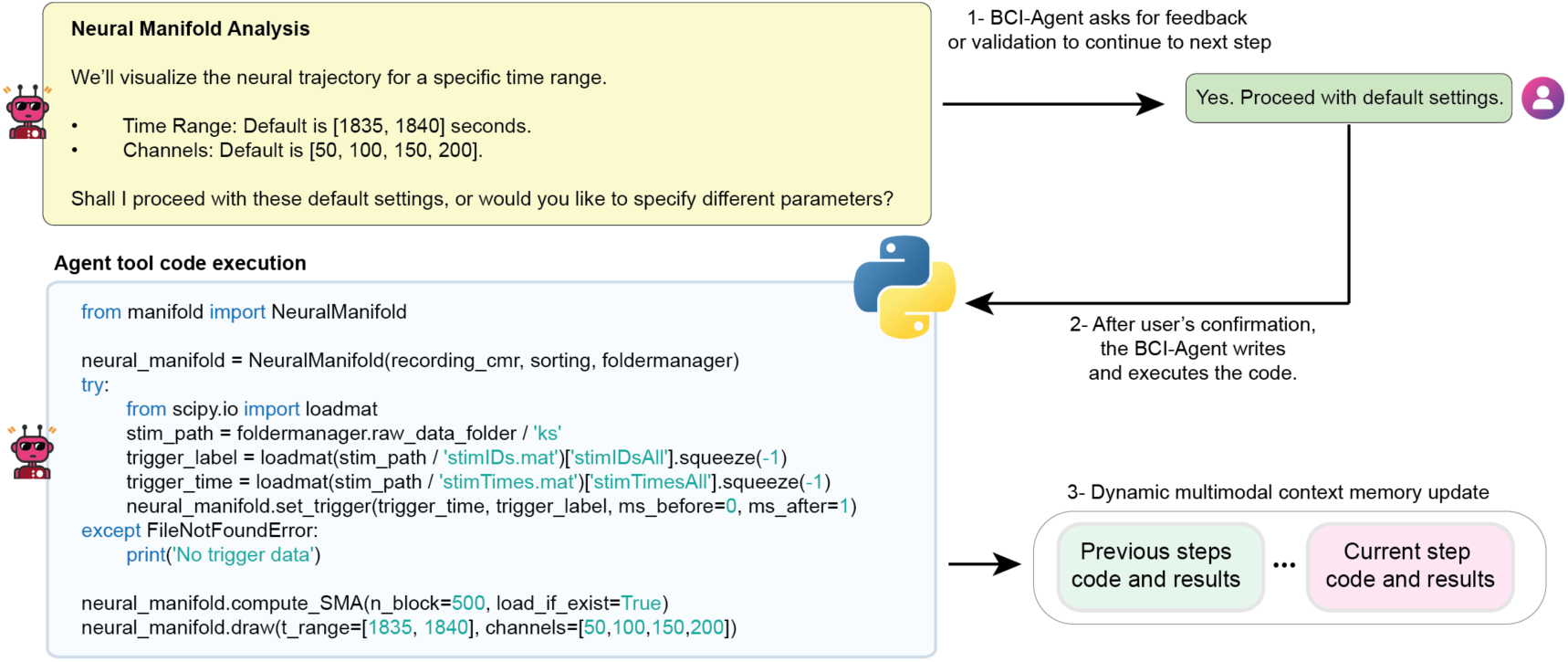
Interactive code execution and multimodal memory update within the BCI-Agent framework. Step-by-step interaction between the user and BCI-Agent during neural manifold analysis. Top left, the agent proposes default settings for visualizing neural trajectories, including a specified time range and selected channels. (1) Top right, the user confirms whether to proceed with the default parameters or customize them. (2) Center, after receiving confirmation, BCI-Agent generates and executes Python code to compute the SMA for manifold visualization. The script imports the required modules, loads trigger data, sets timing parameters, and performs dimensionality reduction with default or user-defined inputs. (3) Bottom right, the results, generated plots, and code from the current step are stored in the agent’s multimodal memory alongside previous steps, enabling dynamic, step-aware context management for downstream reasoning and reproducibility.

**Extended Data Fig. 2.**
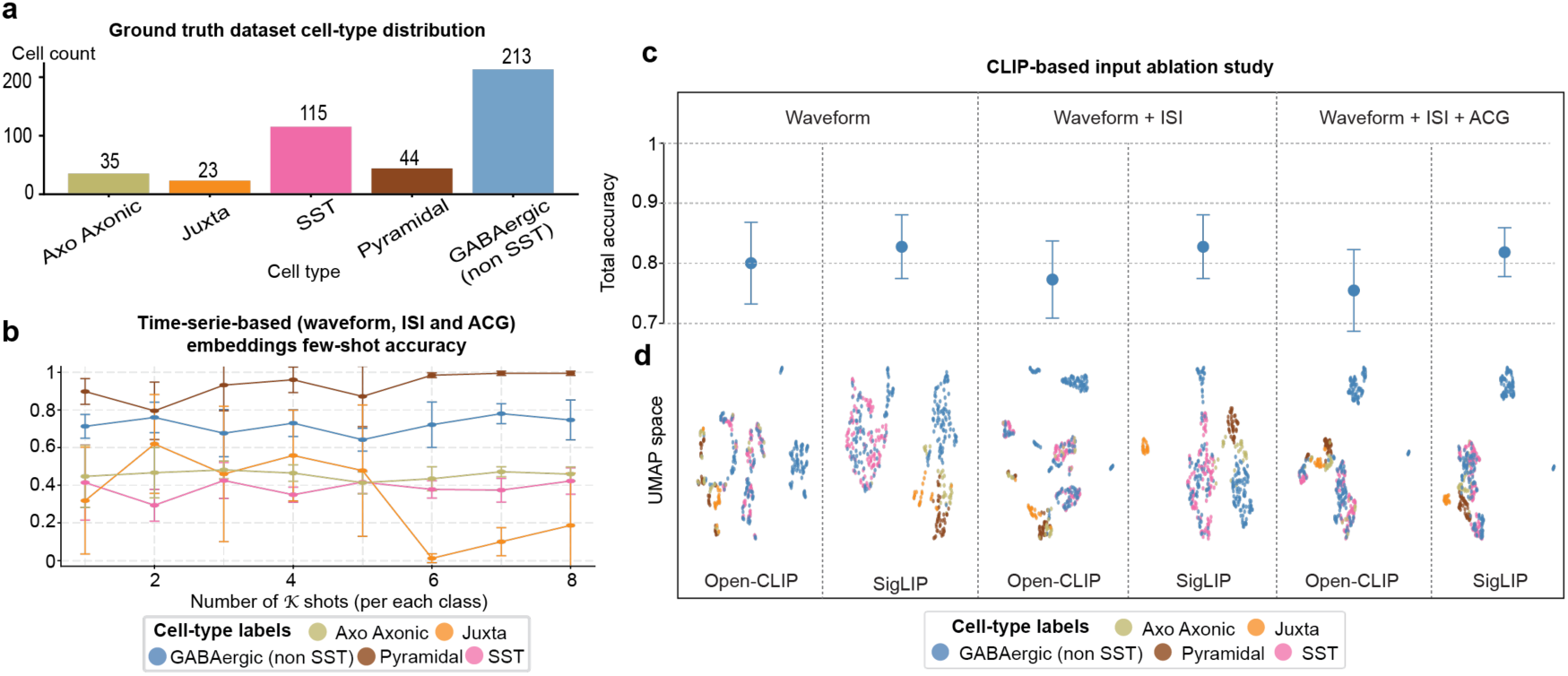
VLM-based classification of neuronal cell types using electrophysiological features. **a**, Ground-truth distribution of cell types in the dataset, including morphologically defined classes (Axo-Axonic, Juxtacellular), excitatory pyramidal neurons, SST-expressing interneurons, and a grouped GABAergic (non-SST) class comprising PV, VIP, VGAT, and PVVGAT. **b**, Few-shot classification accuracy using the waveform, ISI and ACG timeseries concatenated as embeddings across increasing numbers of shots **c**, Classification accuracy for OpenCLIP and SigLIP models across three input feature modalities: waveform only, waveform + ISI, and waveform + ISI + ACG. Accuracy is computed by training a random forest classifier of the generated embeddings after the CLIP visual encoder with 5% of the dataset used for testing. **d**, UMAP visualizations of cell embeddings from each model and modality combination. All results represent the mean (± s.d.) across five random seeds per model.

**Extended Data Fig. 3.**
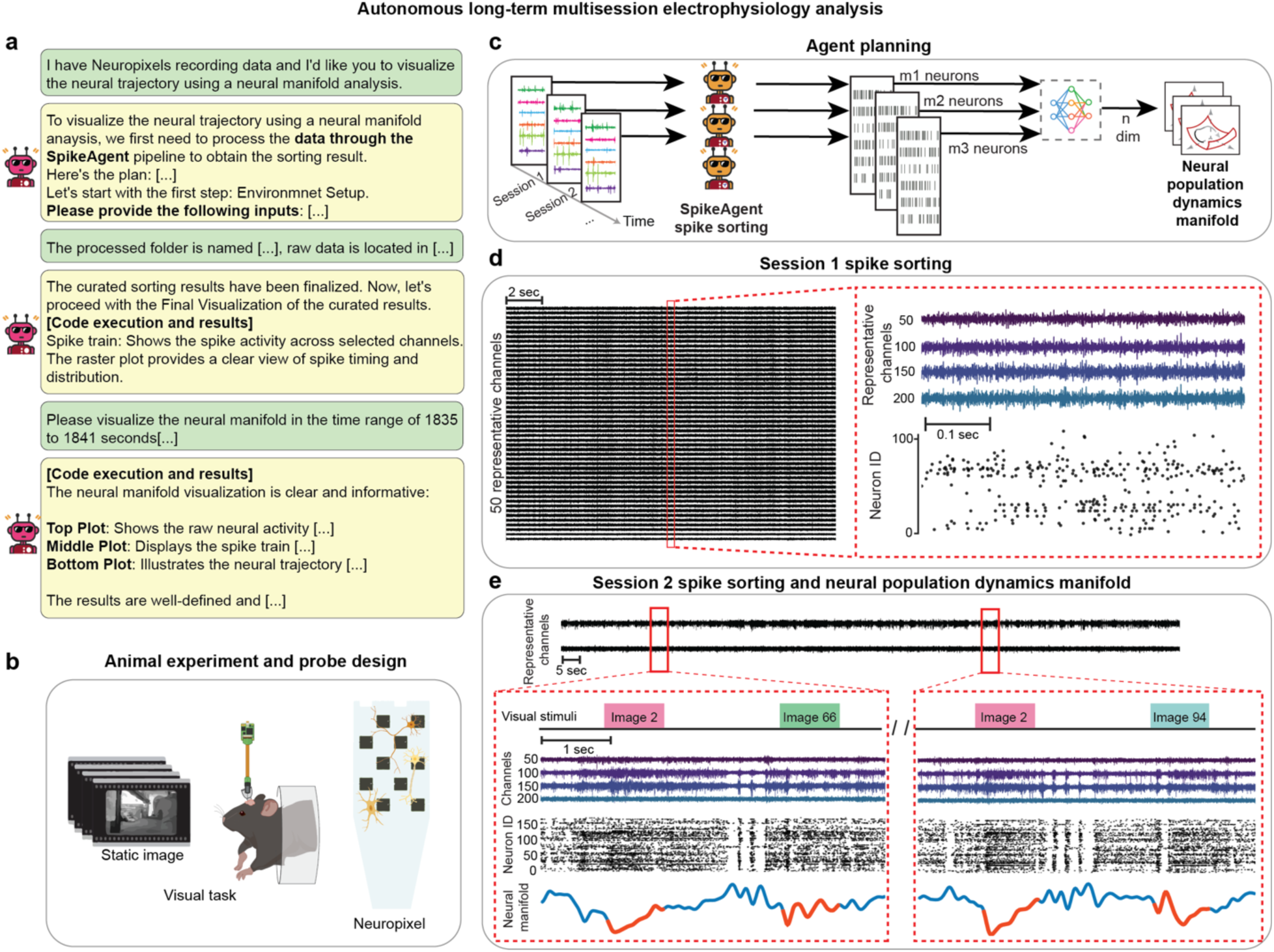
Autonomous, multi-session and aligned spike sorting and neural dynamics analysis in Neuropixels. **a,** Example user-agent conversation for a multi-session analysis workflow. The user provides raw data paths, and the agent initiates session-wise spike sorting and dimensionality reduction to produce neural population dynamics manifolds using the SMA. **b,** Schematic of the experimental setup. Mice perform a visual task while head-fixed, and neural data are recorded using Neuropixels probes. **c,** Agent planning for the multi-session mode. For each recording session, spike sorting is performed using the SpikeAgent tool, and sorted spikes are integrated into population-level trajectories across time. **d,** Spike sorting output for Session 1, showing the raw raster plot from 50 representative channels and example waveform and spike raster visualizations from identified neurons. **e,** Combined spike sorting and population manifold analysis for Session 2 (recording after 56 days of Session 1). Visual stimuli and their alignment with spike rasters and neural manifold trajectories are shown across image-triggered segments.

**Extended Data Fig. 4.**
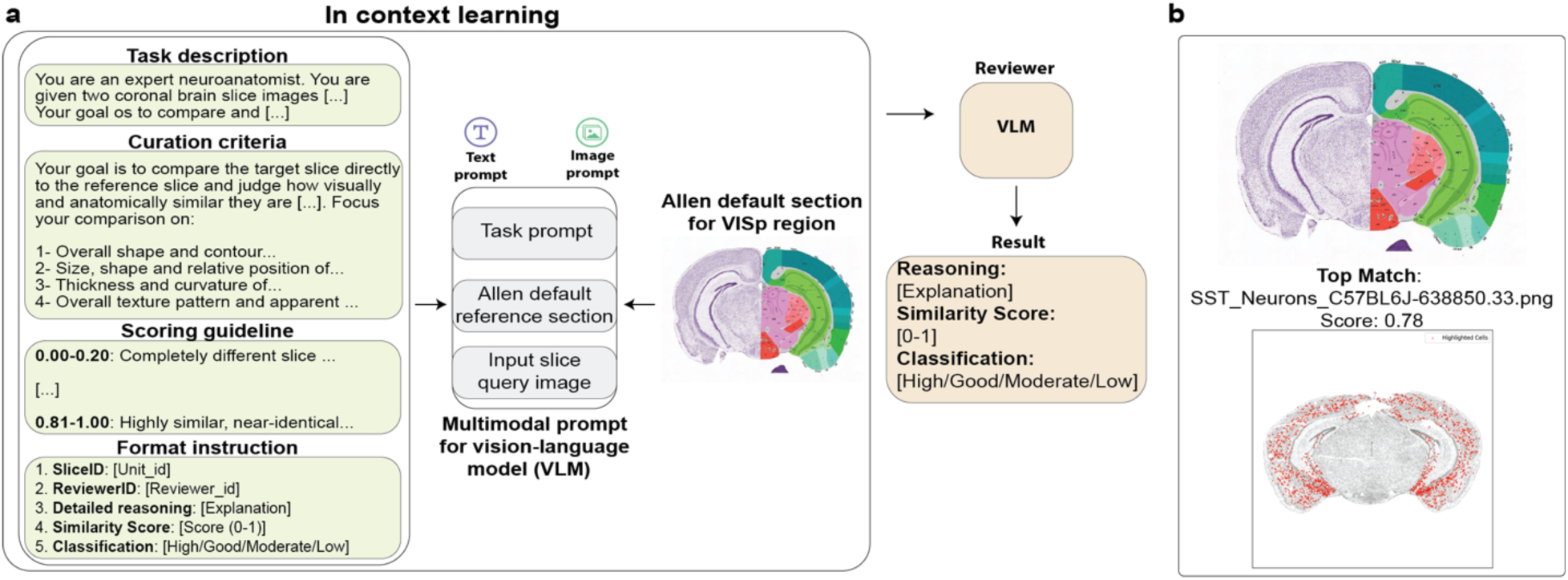
Molecular atlas fact-checking: vision-language models can identify anatomically relevant brain regions using multimodal prompts. **a,** Schematic of the in-context learning setup used to validate anatomical correspondence between experimental input slices and reference atlas sections. A VLM receives a multimodal prompt including a natural language task description and two images: the Allen Institute’s default reference section for the desired region (i.e. VISp) and, one by one, the agent-provided input slices. **b,** Example output showing the top match for the agent-provided SST neuron expression map.

**Extended Data Fig. 5.**
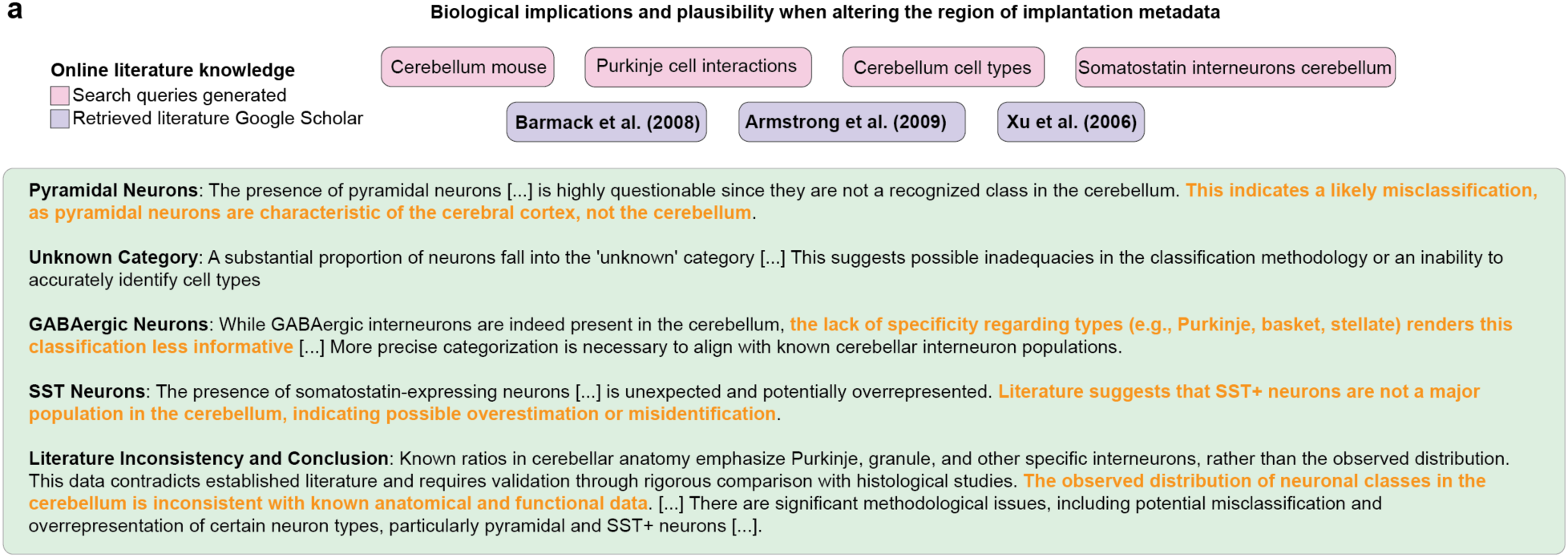
Biological implications and plausibility report with altered region of implantation metadata. **a**, Summary of biological plausibility and consistency assessment comparing inferred human neuronal subtype proportions against established reference data from molecular atlases and literature, detailing alignment and discrepancies between inferred data and known neuronal subtype distributions.

**Extended Data Fig. 6.**
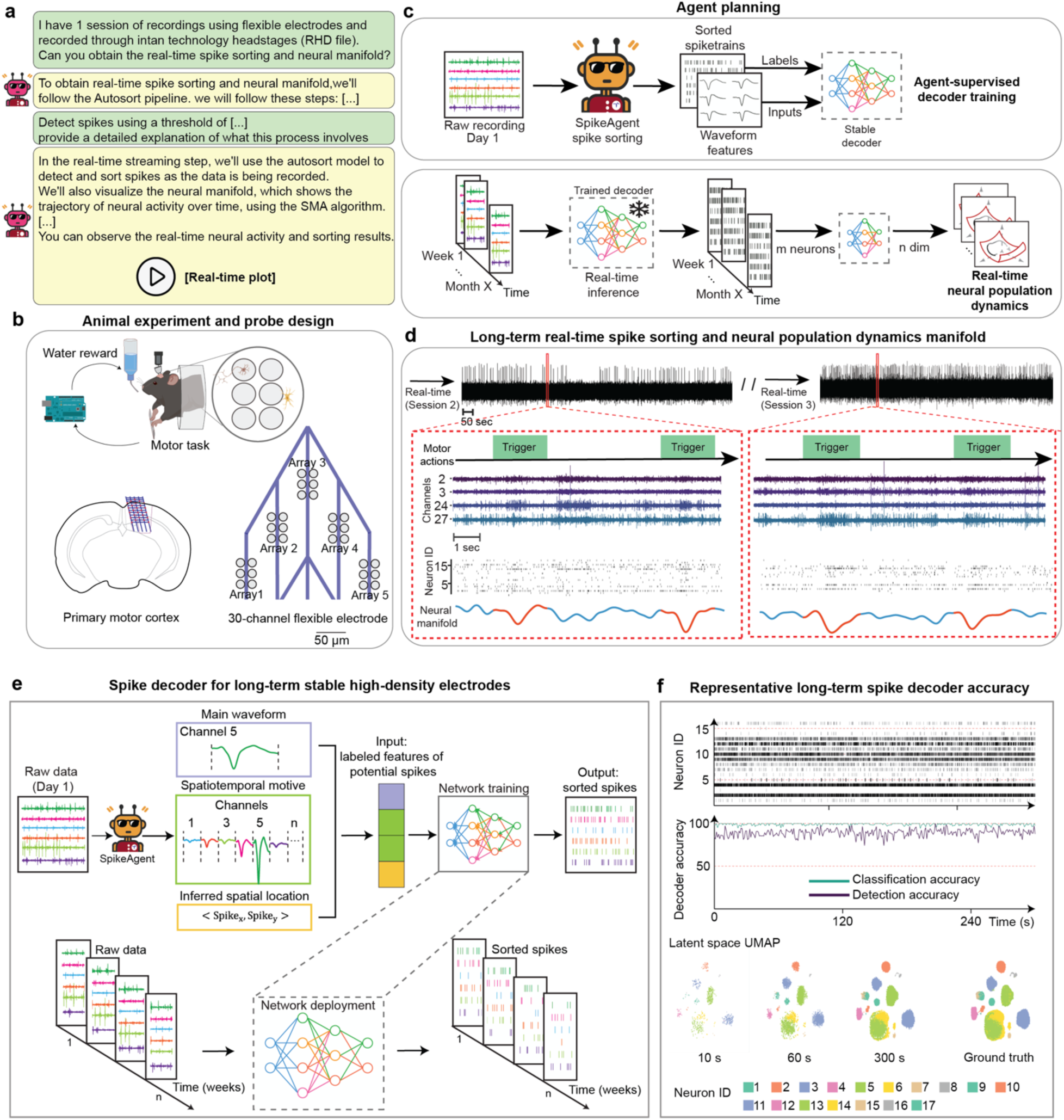
Real-time autonomous, multi-session and aligned spike sorting and neural dynamics analysis in flexible electrodes. **a**, Example user-agent interaction for aligned spike sorting and real-time neural dynamics. The user requests spike sorting and manifold inference on real-time streaming data from a flexible electrode experiment. **b,** Schematic of the motor task experiment. A mouse performs a water-rewarded motor task while recordings are collected from 30-channel flexible electronics implanted in the primary motor cortex. **c,** Real-time agent planning. SpikeAgent first performs spike sorting on an initial reference session to train a neural decoder. The decoder is then deployed in real time on incoming data to classify spikes and infer neural trajectories. **d,** Example output from the real-time mode showing spike activity and neural trajectories aligned to behavioral triggers across multiple weeks. **e**, Overview of the spike decoder architecture used for long-term stable unit tracking. On Day 1, BCI-Agent performs initial spike sorting using the SpikeAgent tool and extracts labeled spike features, including waveform shape, spatiotemporal motifs across channels, and inferred spatial location. These features are used to train a deep learning-based decoder that outputs sorted spikes from raw neural data. Once trained, the decoder is deployed across all subsequent sessions, enabling consistent identification of neuron-specific spike trains over weeks of recording. **f**, Decoder performance during long-term deployment. Top: Example raster plots showing consistent neuron ID assignments over time. Middle: Accuracy of spike detection and classification accuracy across time, indicating stable decoding performance compared to standard spike sorters. Bottom: UMAP visualizations of decoder latent space at various time points (10s, 60s, 300s) and the labeled ground truth distribution.

**Extended Data Fig. 7.**
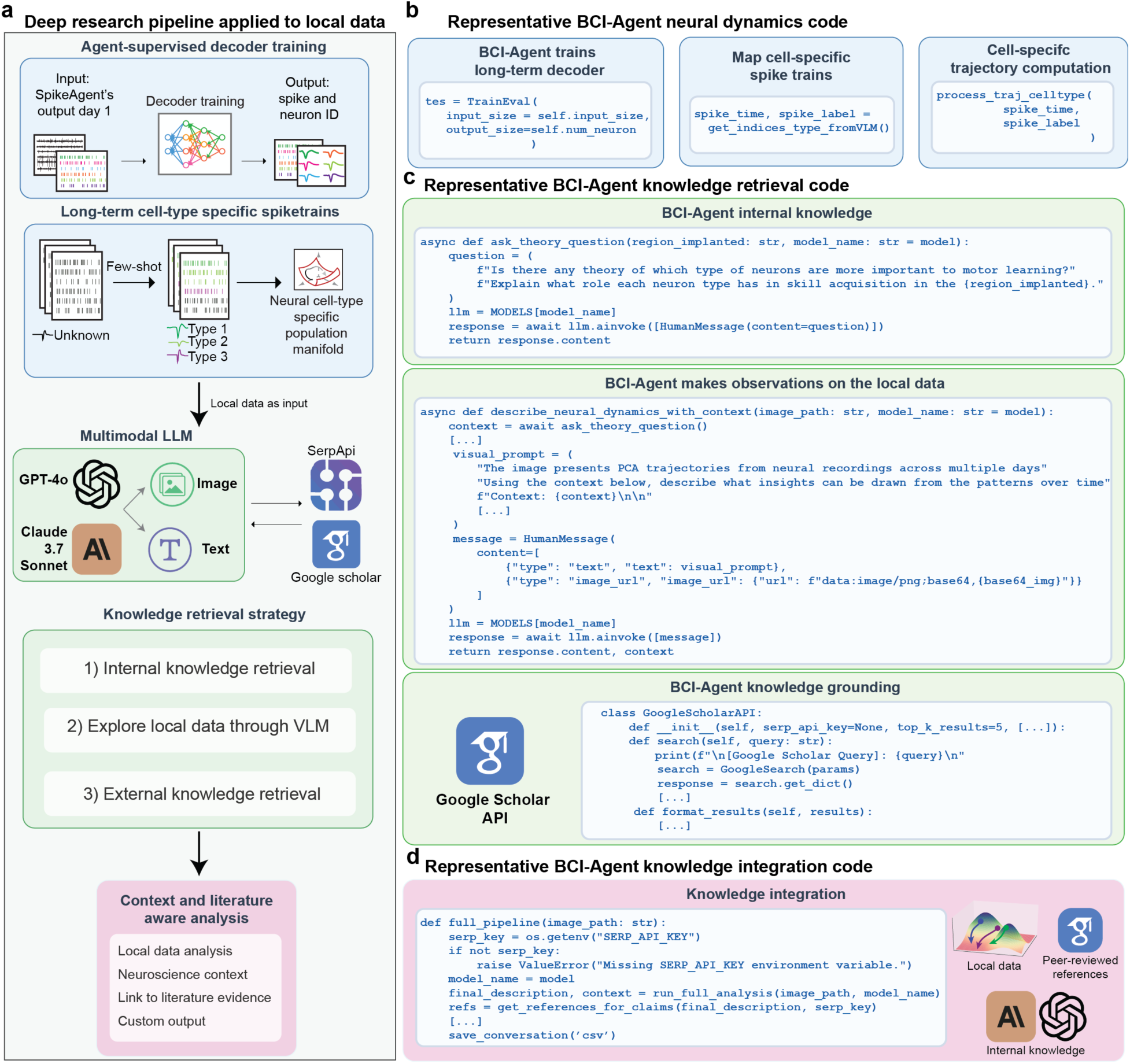
Autonomous knowledge retrieval, integration, and synthesis from local data. **a,** Overview of the BCI-Agent framework for deep research integration with local electrophysiological data. Top: Agent-supervised training of a spike decoder from Day 1 output to learn long-term stable neuron identities and cell-type–specific labels. Bottom: Trained cell-type–specific spike trains are used to build low-dimensional population manifolds, enabling few-shot classification of previously unknown neurons. Local data are analyzed in conjunction with contextual information through a multimodal LLM, integrating visual, textual, and image-based input, and using tools such as SerpAPI^64^ and Google Scholar for external literature access. The agent’s knowledge retrieval strategy consists of three stages: (1) internal retrieval from its LLM; (2) analysis of local neural data using VLM tools; and (3) external retrieval from scientific literature. **b,** Representative code used by the BCI-Agent to train a long-term decoder and compute cell-specific spike trains and trajectories. The spike time and spike label outputs are mapped to neural cell types based on few-shot classification from the VLM. **c,** Code snippets demonstrating the BCI-Agent’s knowledge retrieval capabilities. Top: Internal knowledge retrieval from the agent’s LLM by asking theoretical neuroscience questions (e.g., about region-specific cell-type roles in learning). Middle: BCI-Agent contextualizes local data, describing patterns in neural dynamics via visual prompts based on image representations (e.g., PCA trajectories). Bottom: External grounding through automated literature search using the Google Scholar API. **d,** Example integration code where the agent combines internal, local, and external sources. Final outputs include literature-supported contextual interpretations of cell-type–specific neural dynamics, saved in user-defined formats such as CSV.

**Extended Data Fig. 8.**
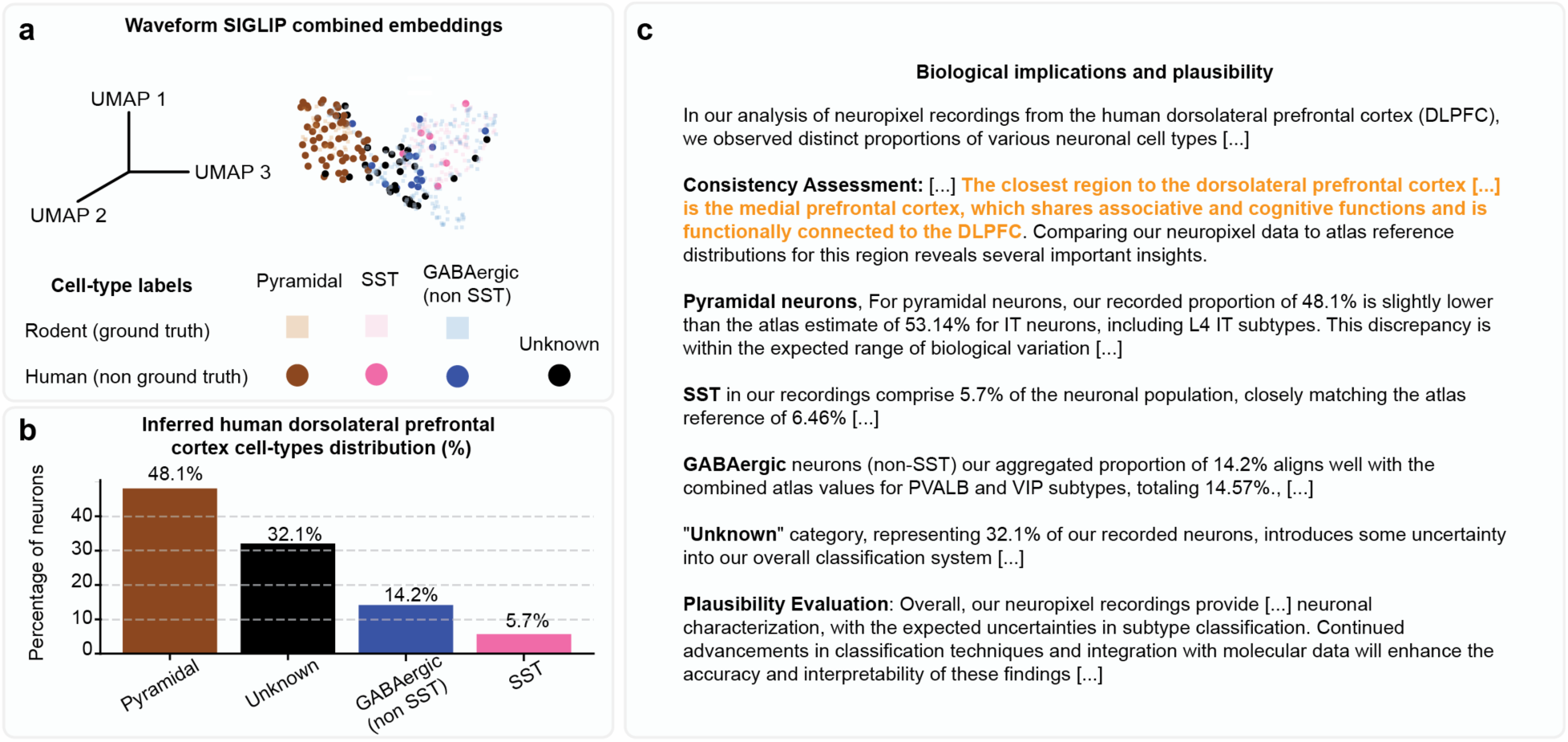
Cross-species transfer of neuronal cell-type inference from rodent to human single-neuron recordings in dorsolateral prefrontal cortex. **a,** UMAP visualization of combined rodent (ground truth) and human (non-ground truth) neuronal waveform embeddings generated by the pretrained CLIP-based model, illustrating the clustering of inferred cell types. **b**, Bar plot summarizing the inferred neuronal cell-type distribution percentages from human Neuropixels recordings, identifying proportions of pyramidal, SST, GABAergic (non-SST), and unknown neuron classes. **c**, Summary of biological plausibility and consistency assessment comparing inferred human neuronal subtype proportions against established reference data from molecular atlases and literature.

